# A theory of physiological similarity in muscle-driven motion

**DOI:** 10.1101/2022.12.13.520057

**Authors:** David Labonte

## Abstract

Muscle contraction is the primary source of all animal movement. I show that the maximum mechanical output of such contractions is determined by a characteristic dimensionless number, the “effective inertia”, Γ, defined by a small set of mechanical, physiological and anatomical properties of the interrogated musculoskeletal complex. Different musculoskeletal systems with equal Γ may be considered *physiologically similar*, in the sense that maximum performance involves equal fractions of the muscle’s maximum strain rate, strain capacity, work and power density. I demonstrate that there exists a unique, “optimal” musculoskeletal anatomy which enables a unit volume of muscle to deliver maximum work and power simultaneously, corresponding to Γ close to unity. External forces truncate the mechanical performance space accessible to muscle by introducing parasitic losses, and subtly alter how musculoskeletal anatomy modulates muscle performance, challenging canonical notions of skeletal force-velocity trade-offs. Γ varies systematically under isogeometric transformations of musculoskeletal systems, a result which yields new fundamental insights into the key determinants of animal locomotor performance across scales.

---

Muscle, the “prime mover” of the animal kingdom, is used for acts of tender kindness, devastating brutality, and astonishing grace. By conversion of chemical into mechanical energy, it enables contractions long-lasting and cyclical, actions fast and forceful, movements precise and reflexive. Whether an animal swims, runs, crawls or flies; whether it is smaller than the tip of a sharp pencil or heavier than 15 school buses; whether it first appeared millions or only thousands of years ago — muscle is what gets it about [1–4].

Juxtaposed to this diversity stands the observation that many functional, physiological and ultrastructural features of muscle vary remarkably little [1, 5, 6], suggesting the existence of general limits to what it can achieve. Identification of these limits, and assessment of their consequences for locomotor performance and musculoskeletal anatomy, has a long history in biomechanics and muscle physiology [e. g. 7–24]. A particularly successful and thus popular approach in comparative studies of animal locomotion has been the notion of similarity indices, derived via dimensional analysis, and introduced to biology in the hope to replicate the success it afforded in the physical sciences [e. g. 11, 13, 14, 25]. Remarkably, the two dominant similarity indices for an-imal movement — the Froude and the Strouhal number — consider elastic and gravitational forces as the agents of motion [3]. The scarcity of similarity theories which make explicit reference to muscle [see e. g. 9, 16, 17, 21, 22] may be partially explained by the complexity of muscle as a motor: muscle force, work and power output depend in non-trivial fashion on muscle strain rate [26], strain [27], and contractile history [28]. Appropriate assessment of muscle performance thus requires coupling these characteristic properties with both internal and external forces, in order to avoid results that are mechanically possible, but physiologically prohibited [29–31]; a task rarely addressed and so challenging that it typically requires numerical resolution [e. g. 15, 19, 32–34].

In this text, I investigate how the interaction between physical constraints, muscle physiology, and the anatomy of musculoskeletal systems places bounds on the mechanical performance space accessible to muscle. I demonstrate analytically that the maximal mechanical output of every muscle contraction is governed by the competition between two distinct limits, which arise from physiological and anatomical constraints. The relative importance of these constraints characterises the degree of ‘physiological similarity’ of muscle contractions, a met-ric defined by a dimensionless number that enables direct comparison of musculoskeletal systems across size, and physiological and anatomical make-up.

## Two speed limits for muscle-driven motion

Consider the seemingly simple example of a muscle with constant gear ratio *G*, capable of exerting a maximum force *F*_*max*_, as it contracts to accelerate an object of mass *m*.^*^ What is the maximum speed it can impart in a single contraction? Mass, maximum force, and gear ratio together determine the net acceleration (*a*), *a* = *F*_*max*_*Gm*^−1^, of dimension length per time squared [L t^-2^]. Identifying a maximum speed *v* of dimension length per time [L t^-1^] thus requires to specify either a time (*t*), *v*_*t*_ ∼ *at*, or a displacement (*δ*), 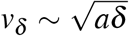. These results may be recognised as the first time- and path-integrals of Newton’s 2^nd^ law, respectively, which link the speed imparted to the delivered impulse (*p*), *mv* ∼ *p* ∼ ∫ *F*_*max*_ *Gdt*, or the work done (*W*), *mv*^2^ ∼ *W* ∼ ∫ *F*_*max*_*Gdδ*. Unfortunately, this analysis does not yet yield the maximum speed the muscle can impart, for Newtonian mechanics alone provides no information about what the appropriate displacement or time might be. Because no muscle can contract in perpetuity, both boundary conditions may be identified through introduction of appropriate physiological and anatomical constraints.

In order to develop some intuition for how such constraints enter the problem, consider the joyful if simplistic analogy of riding a bicycle: If the gear is much too small, it is impossible to increase speed, because the muscle cannot contract quickly enough to accelerate the pedals; the best it can do is to keep them spinning at their current speed. If the gear is much too large, in turn, the muscle will accelerate the pedals rather slowly, so that the speed imparted at the end of a single contraction cycle will be miniscule. Both scenarios are manifestations of two limits to the speed of muscle-driven motion: How fast a muscle can contract, and by how much it can shorten. The axiomatic limit on muscle strain rate, 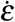, and the anatomical limit on muscle strain, *ε*, can inform the mechanical analysis, for they each impose a distinct limit on the maximum impulse muscle can deliver, and on the maximum work that it can do.

The maximum distance *δ*_*max*_ available for acceleration must be some fraction of the muscle fibre length *l*_*m*_, *δ*_*max*_ = *l*_*m*_*ε*_*max*_*G*^−1^, where I consider the maximum strain *ε*_*max*_ to be independent of the gear ratio. This distance is covered in a time *t*_*max*,_ 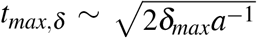 (the factor two arises from integration). Where work and impulse are limited by the strain capacity of muscle, the maximum speed then follows from either maximum distance or time as:

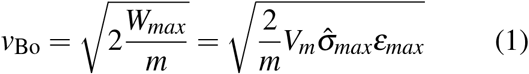

where 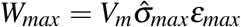 is the maximum work that can be done in a contraction to *ε*_*max*_, *V*_*m*_ is the muscle volume, and 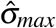 is the maximum *average* stress the muscle can exert as it shortens by *ε*_*max*_.^†^ In recognition of the pioneering work of GA Borelli, who first derived an equivalent result [7], I will refer to eq. 1 as the Borelli-limit to speed, and define the Borelli-number, Bo 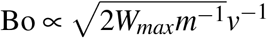.

Consider next the case where muscle work and impulse are limited by the maximum shortening speed of muscle instead. Let this maximum shortening speed be 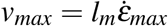, where 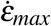 is the maximum strain rate in units of muscle lengths per second. Reaching 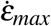 requires accelerating the mass over a distance 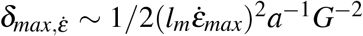, which takes a time 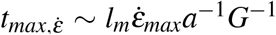. The speed imparted then follows from either as:

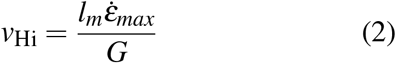

which I will call the Hill-limit to speed, with associated Hill-number, Hi 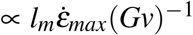, in recognition of AV Hill’s groundbreaking contributions to our understanding of the force-velocity properties of muscle [26]. The Hill-limit may also be derived through a simpler argument: in the absence of series elasticity, the speed of the mass must equal the muscle shortening speed divided by the gear ratio at any time; the mass can thus not move faster than 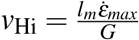.

The Borelli- and Hill-number are dimensionless numbers which may be interpreted as indices of dynamic similarity in muscle-driven motion: where muscle does equal amounts of mass-specific work, movements will have equal Borelli-numbers; equality of Hill-numbers, in turn, implies equal ratios between the muscle contraction speed and the musculoskeletal gear ratio. But which number is appropriate to assess the limit which binds maximum speed?

To answer this question, I once more borrow from the versatile toolbox of dimensional analysis: the ratio between a characteristic speed and a characteristic displacement depends on the acceleration via *v*^2^*δ*^−1^ ∼ *a*. Thus, for vanishing acceleration, the speed gained per unit displacement is negligible, and the muscle will have shortened maximally long before it has reached its maximum shortening speed; the contraction is quasi-static *relative to the maximum strain rate*. If the acceleration is large, in turn, the maximum speed is reached with minimum length change of the muscle, and within miniscule time; the contraction is quasi-instantaneous. Thus, in general, a muscle that contracts against a sufficiently small mass is bound by the Hill-limit (Fig. 1 A & B); the impulse and work it can deliver, and thus the speed it can impart, are limited by its maximum strain rate. In contrast, a muscle that contracts against a sufficiently large mass is Borelli-limited (Fig. 1 A & D); the impulse, work and speed it can deliver are limited by its strain capacity.

**Figure 1.**
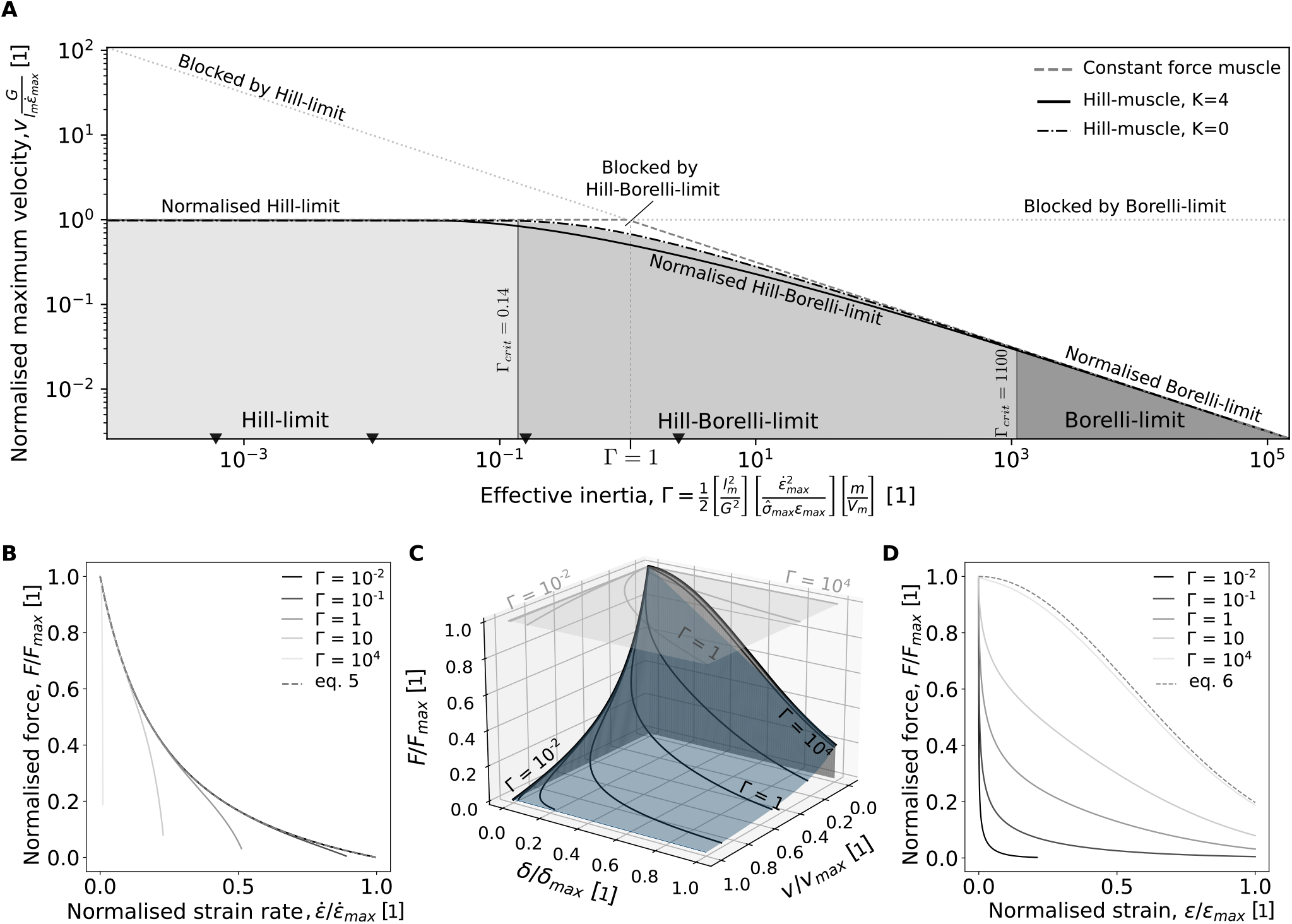
(**A**) The maximum speed muscle can impart in a single contraction depends on the effective inertia, Γ, of the mass-musculoskeletal system. For Γ < 1, the speed is bound by the Hill-limit (eq. 2), which arises because muscle has a finite maximum strain rate, 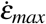. For Γ > 1, the speed is bound by the Borelli-limit (eq. 1), which arises because muscle can only shorten by a fraction of its length, *ε*_*max*_*l*_*m*_. For a constant force muscle, the Hill- and the Borelli-limit are exact expressions for the maximum speed (dark grey dashed line), and the transition between both occurs abruptly at Γ = 1. For a Hill-muscle (black solid & dashed lines), where the force depends both on muscle strain rate and strain, the Hill- and the Borelli-limit are asymptotic, reasonably accurate only below or above a critical value Γ_*crit*_ (see SI); for intermediate values of Γ, the speed is bound by the Hill-Borelli-limit instead, which reflects reductions in work capacity due the strain-rate dependence of the muscle force. Black triangles at the bottom indicate an estimate for the effective inertia of the hindlimb of a generalised tetrapod with a body mass of 10 g, 1 kg, 100 kg, and 1 tonne, respectively (left to right, see text for details). (**B**) The effective inertia controls the fraction of the maximum strain rate muscle can access. For vanishing Γ, the contraction may be considered quasi-instantaneous, and the relation between force and velocity is described solely by the FV relationship of the muscle (cf. to (**D**), and see eq. 5). (**C**) The solutions of all possible equations of motion can be visualised on a 3D landscape which relates the normalised force to the normalised displacement and velocity. For a constant force muscle, the resulting contractile landscape is a horizontal plane (grey, on top), but for a Hill-muscle, it describes a complex 3D envelope (blue), shaped by the projections of the FV and FL relationships, respectively. A muscle with small operates in the Hill-limit and contracts close to the FV plane; a muscle with large Γ operates in the Borelli-limit and contracts close to the FL plane. (**D**) The effective inertia determines the fraction of the maximum strain over which muscle can accelerate. For diverging Γ, the relation between force and displacement approaches the FL relationship (dashed line), and the contraction may thus be considered quasi-static with respect to the maximum relative speed (cf. to (**B**), and eq. 6).

This dimensional argument is simple, but unsatisfying – how large is a sufficiently large, and how small is a sufficiently small mass? To render the qualitative argument quantitative, I next normalise the mass with the relevant physiological properties of muscle, with the aim to find the “effective inertia” of the musculoskeletal system.

### The effective inertia of a muscle contraction

The Borelli- and Hill-limit are both absolute: a muscle that has exhausted its strain capacity cannot increase the speed of the mass further, even if it could contract faster still; a muscle contracting with its maximum strain rate can no longer accelerate the mass, even if it has not yet shortened by the maximum possible amount. The maximum speed muscle can impart is thus determined by whichever of the two speeds is lower (Fig. 1 A), so that the relevant limit can be identified by consideration of the ratio between the Hill- and the Borelli-number:

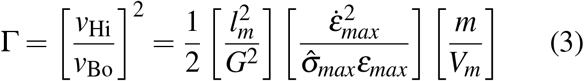

I squared the ratio for convenience, and split the equation in three terms to distinguish between distinct determinants of the effective inertia: the geometry of the musculoskeletal system, represented by the gear ratio and the allocation of a unit volume of muscle into fibre length versus cross-sectional area; the physiology of the muscle, represented by the maximum average stress, strain rate, and strain; and a characteristic density, *m/V*_*m*_, which may be interpreted as a relative investment of muscle tissue compared to the mass that is to be moved. If Γ < 1, the muscle is Hill-limited, and if Γ > 1, the muscle is Borelli-limited; Γ = 1 corresponds to the unique special case for which a constant force muscle operates simultaneously at both limits.

I note that the specific parameter combination in eq. 3 also falls out of more formal dimensional analyses of musculoskeletal systems as a dimensionless mass [see SI in 33, 36, and [16] for a similar result for series elasticity]. Throughout this work, I will discuss several possible interpretations of Γ. For example, Γ also follows as the ratio between the kinetic energy associated with a contraction at maximum strain rate, and the work done in a contraction to maximum strain (see eq. S6 in SI). For Γ < 1, the muscle uses a fraction *ε* = Γ*ε*_*max*_ of its strain capacity to reach its maximum strain rate (Fig. 1 A & B). For Γ > 1, in turn, the muscle uses its entire strain capacity to accelerate to a fraction 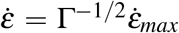 of its maximum strain rate (Fig. 1 A & D). Dimensionless numbers such as Γ can always be interpreted in a number of ways, and I submit that the effective inertia may best be thought of as a *physiological similarity index*, a notion which combines all possible interpretations into one term, and on which I will expand on further below.

It is of obvious interest to estimate the effective inertia of ‘real’ muscle and musculoskeletal systems. In the supplementary information, I derive two such estimates using published data [3, 37–40]. First, for a representative vertebrate muscle contracting against itself, 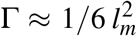 meters^-2^. Thus, even for a long muscle with a fibre length of 1 m, Γ ≈ 1/6 < 1; for a fibre length of 1cm, the effective inertia drops to Γ ≈ 1/6·10^−5^. Second, for a generalised tetrapod hindlimb contracting to accelerate the body mass, Γ ≈ 1/100 *m*^0.6^ kg^-0.6^. Thus, for a shrew with a body mass of 0.01 kg, Γ ≈ 0.6·10^−^3; for a mole with a body mass of 0.1 kg, Γ ≈ 0.0025; for a cat with a body mass of 1 kg, Γ ≈ 0.01; for a dwarf crocodile with a body mass of 10 kg, Γ ≈ 0.04; for a caribou of with a body mass of 100 kg, Γ ≈ 0.16; for a rhinoceros with a body mass of 1000 kg, Γ ≈ 0.63; and for an elephant with body mass of 10,000 kg, Γ ≈ 2.5. The strong size-dependence of Γ is evident, and will be discussed in more detail below.

### The equations of motion landscape and the Hill-Borelli transition for real muscle

The above analysis is an exact result for a muscle which generates a constant force throughout the contraction. In reality, however, muscle is more complex a motor, and the force it generates is a function of both its relative strain rate, 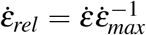, and its rel-ative length, 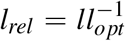 [26, 27]: muscle produces maximum force during an isometric contraction, 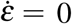, at “optimal” length, *l* = *l*_*opt*_, and less force for any deviation from these conditions. Unfortunately, as the muscle contracts to accelerate the mass, it must change both its length and contractile speed; the net force it generates — and thus the acceleration the mass experiences — consequently varies continuously throughout the contraction. In order to assess the effect of this dynamic complexity, I now introduce the force-length (FL) and force-velocity (FV) properties of muscle. I make three simplifying assumptions to aid this analysis: the muscle activation time constant, *t*_*act*_, is much smaller than the characteristic acceleration time, 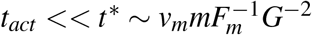, so that the muscle can be considered fully activated throughout; I consider a single contraction from rest at the optimal length, so that the relative length can be re-expressed via the strain, 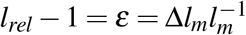; and the normalised FV and FL functions, 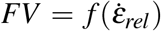 and *FL* = *f* (*ε*), respectively, are independent, and each modulate the generated muscle force such that it is equal to some fraction of its maximum value (dashed lines in Fig. 1 B & D). The muscle force may then be written as 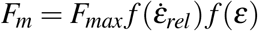.

In order to develop some intuition for the effect of FV and FL properties on muscle contraction dynamics, I briefly return to the simpler case of a muscle which generates a constant force throughout the contraction, i. e. 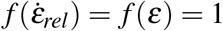, and thus *F*_*m*_ = *F*_*max*_. The contraction dynamics — that is the change of speed and displacement with time — are governed by the equation of motion (EoM), *a* = *F*_*max*_*Gm*^−1^. All possible solutions of this EoM can be visualised I a single plot of the normalised net force against the normalised displacement and velocity, respectively (Fig. 1 C). Because the net force is constant throughout the contraction, all such solutions lie in a horizontal plane (grey plane at the top of Fig. 1 C). Consider now the path inscribed onto this plane by contractions with different effective inertias; as illustrative example, let the very same muscle contract against different masses. For any specific mass, the contraction describes a unique trajectory onto the EoM plane: If the mass is very small, Γ vanishes and the mass thus gains normalised speed rapidly, and with minimum normalised displacement; the contractile trajectory is almost parallel to the FV-line. If the mass is large, in turn, Γ diverges and the normalised speed remains small throughout the displacement; the contractile trajectory is almost parallel to the FL-line. For intermediate masses, corresponding to Γ close to unity, the contraction involves significant changes in both normalised speed and displacement, and thus describes a more complex FVL trajectory (Fig. 1 C).

Consider next the more complex case of a Hill-muscle with FV and FL properties, governed by the EoM, 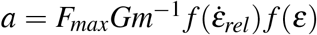. The two horizon-tal lines which defined the EoM plane for a constant-force muscle are now curves, and the EoM plane is thus transformed into a three-dimensional EoM landscape, shaped by their projection (Fig. 1 C). The principal logic and interpretation, however, remain the same: If the muscle contracts against a vanishing mass, Γ is small, the acceleration is large, and the muscle reaches its maximum velocity with a small change in length. The EoM of the contraction is defined solely by the muscle’s FV properties, *f* (*ε*) ≈ 1, so that the contractile trajectory is approximately equal to the FV-curve (Fig. 1 B-C). If the muscle contracts against a diverging mass, in turn, Γ is large and the acceleration is small, and the muscle will have shortened by its maximum amount long before it has reached its maximum strain rate. The EoM of the contraction is defined solely by the muscle’s FL properties, 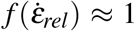, so that the contractile trajectory is approximately equal to the FL-curve (Fig. 1 C-D). The relative FV and FL functions may thus be interpreted as the sole contributors to the EoMs of two contractions which represent unphysical extremes: a quasi-instantaneous contraction which reaches the maximum strain rate with vanishing strain; and a quasi-static contraction, during which a muscle shortens by its maximum strain with a vanishing increase in strain rate.

Although the introduction of FV and FL-properties leaves the effect of variations in Γ qualitatively unaltered, there exists a material difference. For a constant-force muscle, the Hill- and the Borelli-limit are exact expressions for the maximum speed muscle can impart; the transition between both limits is sharp, and occurs at Γ = 1, where the muscle reaches its maximum contraction velocity exactly when it has contracted to the maximum strain (Fig. 1 C). For a Hill-muscle, the Hill-limit remains intact, but the Borelli-limit is now only exact in the limit of diverging Γ, for which the contraction becomes quasistatic; the imparted speed is smaller than *v*_Bo_ for any other contraction to *ε*_*max*_. This result may be understood qualitatively as follows: the Borelli-limit is the speed that can be imparted during a contraction which involves the maximum possible work output, or, equivalently, the maximal possible area underneath the FL-trajectory. For a constant-force muscle, this area is maximised for *any* contraction to *ε*_*max*_. For a Hill-muscle, however, the area is maximal only for a quasi-static contraction; any increase in muscle strain rate reduces the muscle work output compared to this maximum [Fig. 1 D and ref. 19]. Because the work output during realistic displacement-limited contractions is influenced by both the FL and the FV-function, I will refer to the corresponding speed limit as the Hill-Borelli-limit, *v*_Hi-Bo_.

Evaluation of the Hill-Borelli-limit requires solution of the EoM 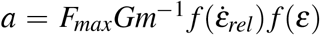. This EoM is a non-linear differential equation, and the chance of a closed-form solution is maximised, if still slim, if it is re-written as an integral over the path instead of the time, which permits separation of variables (see SI for detailed derivations and further discussion of the results which follow):

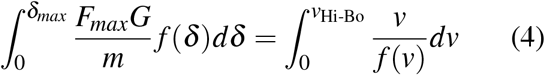

where I used the coupling *δ* = *l*_*m*_*εG*^−1^ and 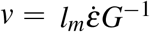, respectively, to re-express the strain and strain rate in terms of the integration variables. Equation 4 may be recognised as the Work-Energy-Theorem, which relates the work done by the muscle to the resulting change in the kinetic energy.

Up to this point, the exposition is agnostic to the form of the FL and FV relationship, and thus general. Further evaluation however requires a specific choice, which is not trivial, for there are no accepted first principle forms of the FL and the FV functions of striated muscle [but see 41, for exciting recent developments]. I proceed with two common forms, not to proclaim their superiority, but merely by way of example; the procedure laid out here may be followed just as well for any other choice. I describe the FV relationship with a normalised Hill-relation [1, 26, see dashed line in Fig. 1 B]:

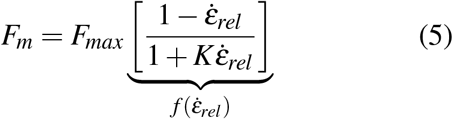

Here, *K* is a dimensionless constant of order unity, which controls the curvature of the FV-relationship [1]. The FL-relationship, in turn, may be written as [e. g. 42, see dashed line in Fig. 1 D]:

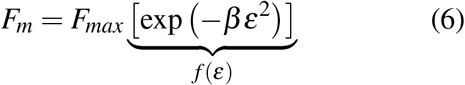

where *β* is a dimensionless shape parameter, which controls how quickly the force decays with strain.

The two integrals in eq. 4 have a closed-form solution for these two forms, but an explicit writing in terms of *v*_Hi-Bo_ is only possible for a linear FV-relationship, i. e. *K* = 0 (see SI):

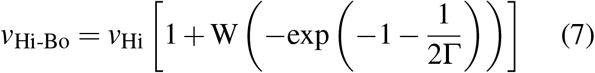

where W is the Lambert W function. Although only exact for the special case *K* = 0, this solution captures all relevant physical (dimensional) parameters, and confirms that the effective inertia remains the key dimensionless number which governs muscle performance: For Γ→ 0, *v*_Hi-Bo_ → *v*_Hi_, and for Γ→ ∞, *v*_Hi-Bo_ → *v*_Bo_ (Fig. 1 A. In the SI, I derive limiting values Γ_*crit*_, below or above which the Hill- and the Borelli-number are within 1% of *v*_Hi-Bo_, and may thus be considered reasonable approximations). I thus define the Hill-Borelli-number, Hi-Bo∝ *v*_Hi-Bo_*v*^−1^, as dynamic similarity index which characterises the maximum output of a Hill-muscle across a broad range of Γ. It is instructive to compare the prediction via eq. 7 to (i) the result for a constant force muscle; and (ii) to a numerical result for *K* = 4 and *β* = 6.5 — reasonable values for animal muscle [3, 43, and SI]. *v*_Hi-Bo_ remains within 30% of either result for any value of Γ, and may thus be considered sufficiently accurate unless all experimental quantities are known with small error (Fig. 1 A). I nevertheless proceed with the general form of the Hill-relation in all that follows to retain generality.

Finding an expression for the maximum speed muscle can impart is, at first glance, a simple problem in Newtonian mechanics. But the simplicity of the governing equations is deceiving. No muscle can contract by more than *ε*_*max*_, or faster than with 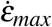. Because these boundary conditions compete and dynamically couple to the mass that is accelerated, and because muscle is a rather peculiar motor with complex FV and FL functions, the problem acquires considerable subtlety. I have demonstrated that the resulting complexity is suitably captured by a single characteristic dimensionless number: the effective inertia, or the physiological similarity index, Γ.

### The effective work and power density of muscle, and the “optimal” geometry of musculoskeletal systems

Equipped with a first order understanding of the relevance of the effective inertia in muscle contractions, we may next interrogate musculoskeletal “design”. A classic concept in the analysis of muscle performance is the notion of a characteristic work and power density; each unit mass of muscle can at most deliver an ostensibly fixed maximum amount of work and power, and these maxima represent putatively suitable metrics to characterise muscle performance limits [4, 7, 9, 10, 22, 33, 36]. What determines whether a mass-musculoskeletal system operates close to these limits?

By definition, muscle operates with maximum work density, *W*_*ρ,max*_, in the Borelli-limit, where it contracts over the largest possible distance with the smallest possible increase in strain rate [Fig. 2 A and 3]; *W*_*ρ,max*_ is a sole function of the muscle’s FL-properties, the muscle density, *ρ*, and the maximum strain, 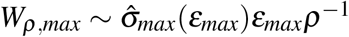 (see eqs. S12 & S16 in the SI). As may be expected by analogy, muscle operates with maximum average power density in the Hill-limit, where it reaches its maximum shortening speed with a minimum loss in force due to length changes (Fig. 2 B); *P*_*ρ,max*_ is a sole function of the muscle’s FV-properties, its density, and a characteristic strain rate, 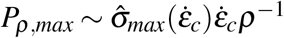(see eq. S13 & S20, and note that the stress is now time-averaged). For a muscle which generates constant force, maximum average power is delivered in contractions to 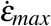, but for a muscle with FV-properties, average power is maximised when the contraction is terminated at a lower strain rate [about half of 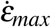for *K* = 4; see SI for a detailed calculation and 44, 45, for related results].^‡^

**Figure 2.**
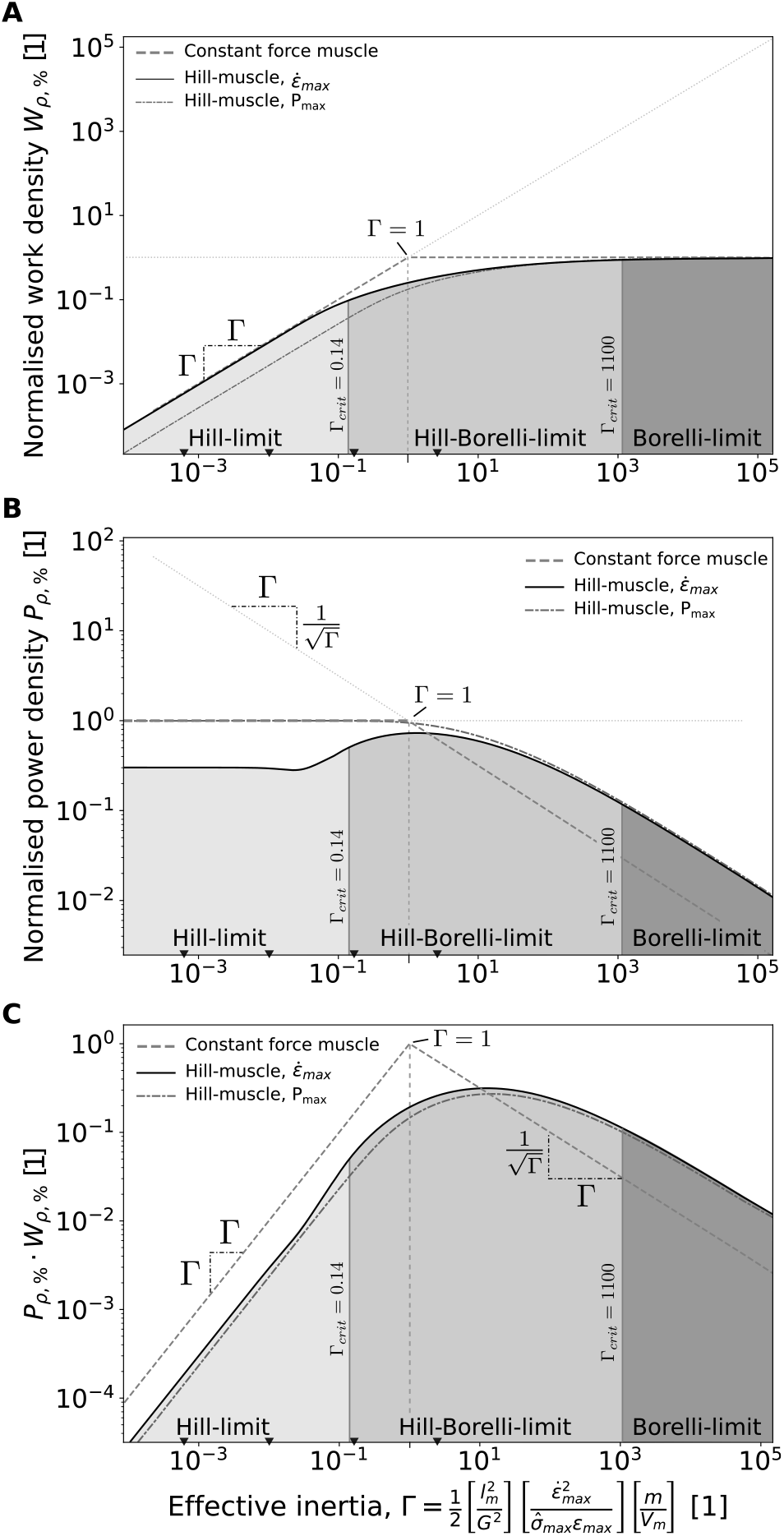
A muscle delivers its maximum work den-sity, *W*_*ρ,max*_, in the Borelli-limit (**A**), and its maximum power density, *P*_*ρ,max*_, in the Hill-limit (**B**). The effective inertia Γ is directly related to the fractions *W*_*ρ*,%_ and *P*_*ρ*,%_ delivered in the Hill- and the Borelli-limit, respectively, as indicated by the slopes. (**C**) Maximising the product between *W*_*ρ*,%_ and *P*_*ρ*,%_ may be considered a design objective: A constant force muscle operating at Γ = 1 can deliver both its maximum work and power density simultaneously. For a Hill-muscle, the maximum product is less than unity and occurs at Γ > 1. Dark grey lines show contractions which result in max-imum average power output, *P*_*max*_, as opposed to close to maximum strain rate, 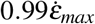 (see text and SI). Black triangles at the bottom indicate an estimate for the effective inertia of the hindlimb of a generalised tetrapod with a body mass of 10 g, 1 kg, 100 kg and 1 tonne, respectively (left to right).

To explore the implications of these observations for musculoskeletal design, I again return first to the simpler case of a constant force muscle: A unit volume of muscle operates with *W*_*ρ,max*_ in the Borelli-limit, and with *P*_*ρ,max*_ in the Hill-limit. It can thus only operate with both *W*_*ρ,max*_ and *P*_*ρ,max*_ if Γ = 1; the unique effective inertia at which the musculoskeletal system is simultaneously in both limits (Fig. 2 C). For Γ < 1, muscle only delivers a fraction of its work density, equal to *W*_*ρ*,%_ = Γ; and for Γ > 1, it delivers only a fraction of its power density, equal to *P*_*ρ*,%_ = Γ^−1/2^ (see SI, and Fig. 2 A-C). The product of the relative power and work densities thus takes a maximum value of unity at Γ= 1, and is equal to Γfor Γ< 1, and to Γ^−1/2^ for Γ > 1. Somewhat fortuitously, Γcan thus also be interpreted as the fraction of the maximum work and average power density muscle can deliver. Indeed, Γ may also be derived as the ratio between the average power and work density of the contracting muscle, 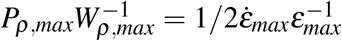, which defines a characteristic physiological time scale; normalisation of this time scale with the time it takes to accelerate to the maximum possible strain rate, 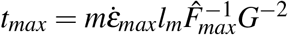, yields the effective inertia.

For a muscle with FV and FL properties, these results are no longer exact, but the implicit scaling relations hold in the limit of small and large Γ, i. e. where the Hill-Borelli-limit is close to the Hill- and the Borelli-limit, respectively (Fig. 2 A-C). Although there no longer exists an effective inertia at which the muscle is simultaneously in both the Hill- and the Borelli-limit, an equivalent optimum may still be defined as the value of Γ which maximises the product between *W*_*ρ*,%_ and *P*_*ρ*,%_. This maximum product is now smaller than unity, and occurs at an effective inertia larger than unity (Fig. 2 C). An exploration of the exact value of Γ which corresponds to this putative optimum as a function of the FL and FV properties of muscle is beyond the scope of this work. However, the implication is clear: operation with small or large Γ means operation with suboptimal work or power density, respectively (Fig. 2 A-C).

It is instructive to note that although neither *W*_*ρ,max*_ nor *P*_*ρ,max*_ depend on the geometry of the musculoskeletal system, nor on the mass muscle contracts against, the ability of muscle to deliver either depends on both (see Fig. 2 A-B and eq. 3). In the classic literature, the influence of mass on the ability of muscle to deliver *W*_*ρ,max*_ or *P*_*ρ,max*_ has rarely been explicitly considered [but see e. g. 19–22, 24, 46], and variations in musculoskeletal anatomy are typically interpreted as controlling a trade-off between force and velocity. The above result demonstrates that this popular interpretation is correct only if muscle operates in the Hill-limit (i. e. Γ < 1), where musculoskeletal geometry determines the extent to which a unit power is split into force versus velocity (see eq. 2). In sharp contrast, in the Borelli-limit (i. e.Γ > 1), the maximum velocity is independent of the musculoskeletal geometry (see eq. 1), which instead merely determines the extent to which a unit amount of work is split into force and displacement, respectively; changing the gear ratio in the Borelli-limit changes the force, but leaves the maximum velocity unaffected. Consider as illustrative example a muscle operating close to an effective inertia of unity: Increasing the gear ratio results in a reduction of both the maximum achievable speed, and the time it takes to reach this maximum speed; the power remains constant but the work done decreases. Decreasing the gear ratio, in turn, leaves the maximum speed unchanged, but increases the time it takes to reach it; the work remains constant but the power decreases [see 20, 23, for other results indicating a workpower tradeoff modulated by musculoskeletal geometry]. At an effective inertia of unity, the musculoskeletal system imparts the maximum possible speed in the shortest possible time.

On the basis of these remarks, it is tempting to consider an effective inertia close to unity as the “optimal design” for a musculoskeletal system. This may well be so, but the notion of a maximum speed, power and work density as sole design objective is likely overly simplistic for at least two reasons. First, where muscle is expected to move a range of masses in stereotypical scenarios, muscle performance is not suitably characterised by a single-valued effective inertia, and will thus deviate by necessity from any putative optimum. Second, delivering the same amount of muscle-mass specific work or power in a different time may well be associated with a variation in efficiency, i. e. the amount of metabolic energy required to deliver a unit amount of mechanical energy. Such variation may then favour effective inertias smaller or larger than unity.

### Parasitic forces and the truncation of the EoM landscape

The above analysis is vulnerable to the reasonable criticism that it is concerned with a case of seemingly limited practical relevance – muscle is never the sole contributor to the net force that accelerates the mass. I will next demonstrate that the initial development, though restricted to a special case, constructed an analytical framework strong enough to carry the burden of further complexity.

The net force is the vector sum of all external forces. Where such forces oppose the muscle force, they will influence muscle dynamics in two distinct ways, distinguished in their generality by their dependence on the FV and FL relationships. Let a muscle contract against an opposing force *P*. The opposing force is “parasitic”, in the sense that it does negative work; it consumes part of the work done by muscle, and in doing so redirects it from kinetic energy to other forms of energy – heat, gravitational potential energy, what have you. To evaluate the partitioning of muscle work into kinetic versus parasitic energy, I derive from conversation of energy:

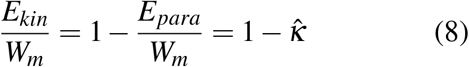

where *W*_*m*_ is the work done by muscle. The reduced parasitic energy 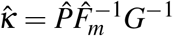 is the fraction of muscle work consumed by the parasitic force, with the immediate consequence that 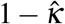 represents the fraction of muscle work which flows into kinetic en-ergy. Because 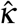 is the ratio of the work done by the parasitic and the driving force, which both act over the same displacement, it is, in general, a ratio of *average* forces. For 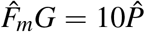, 90% of the work done by muscle flows into kinetic energy, and for 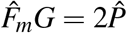 it is half. For 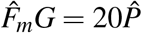, the error in neglecting the parasitic force in the energy balance is less than 5%, and for 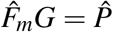, muscle can do zero work; no acceleration is possible at all, and the system is in equilibrium. The reduced parasitic energy can be related to familiar dimensionless numbers, such as the Froude, Strouhal or Reynolds number for gravitational, elastic or viscous parasitic forces, re-spectively, via 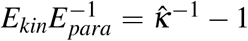 (note that these then differ from their definition in Alexander’s sense, as gravitational and elastic energy are no longer the source of kinetic energy, but a sink for muscle work). I submit that their definition via a ratio of forces is preferable in problems of muscle dynamics, because it places the focus on the agent which caused the change in energy: the contracting muscle.

For a constant force muscle, 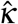 fully captures the influence of parasitic forces on muscle performance. The Hill-number remains unchanged, but the Borelli-number and thus the effective inertia gain an additional term, defined by eq. 8. The net effect of the parasitic force is thus an increase of the effective inertia by a factor 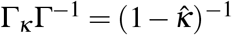; as *P* approaches *F*_*m*_*G*, the constant force muscle is increasingly likely to be Borelli-limited (see Fig. 3 A), and reaching a unit speed takes longer. The interpretations developed above remain valid, but now refer to the work that flows into kinetic energy, rather than the total muscle work.

**Figure 3.**
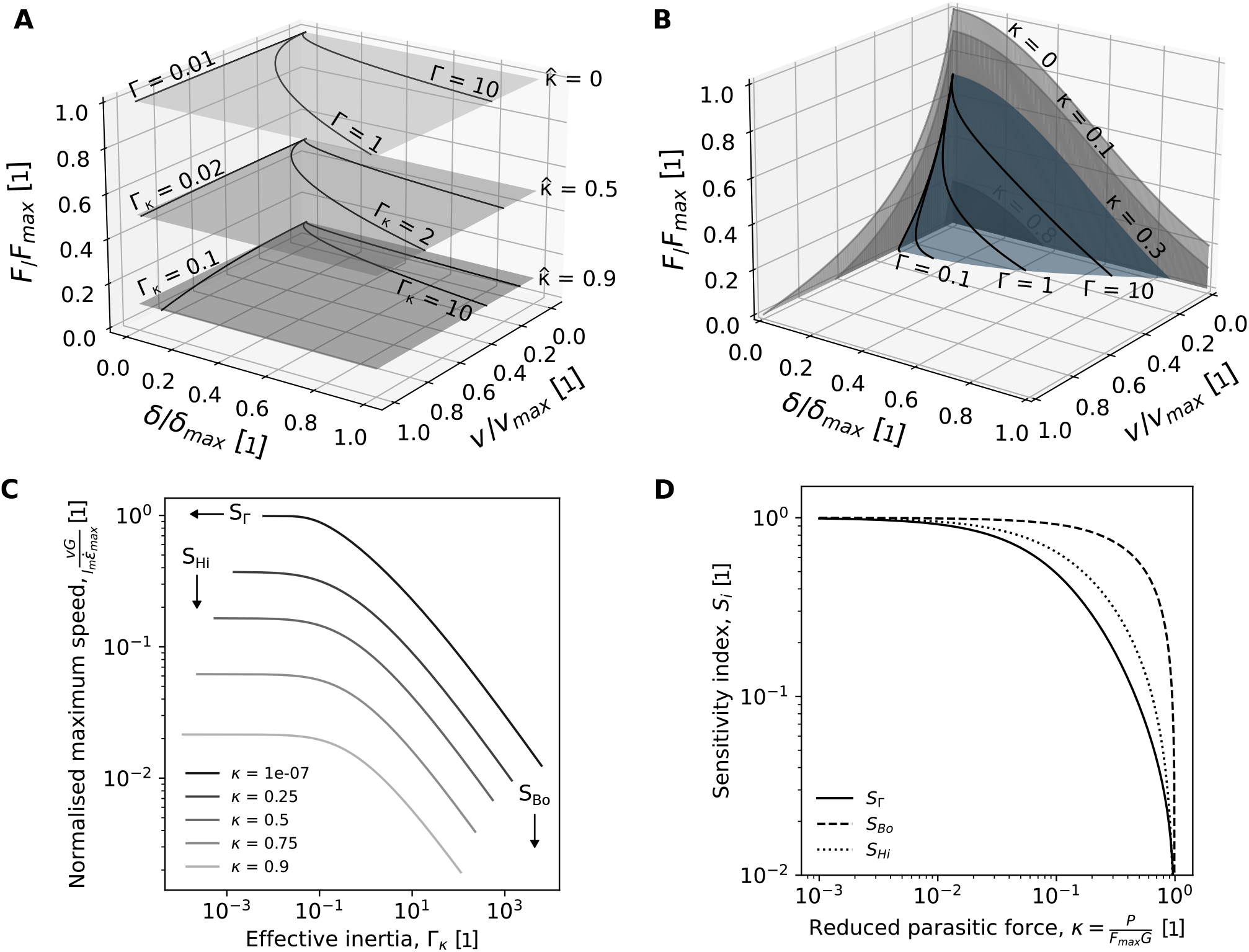
External forces which oppose motion influence muscle dynamics in two distinct ways, distinguished in their generality by their dependence on the force-length (FL) and force-velocity (FV) relationship. (**A**) A constant opposing force, *P*, manifests itself via a downward shift of the equation-of-motion (EoM) plane. *P* consumes part of the work done by muscle and may thus be considered ‘parasitic’. The amount of muscle work which flows into parasitic instead of kinetic energy is quantified by the reduced parasitic energy, 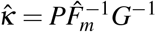, where 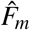 is the muscle force averaged over the displacement, and *G* is the gear ratio. The effective inertia increases with *κ*, as illustrated by the contractile trajectories on each EoM plane. (**B**) For a Hill-muscle, the influence of parasitic forces is more complex. Because the FL and FV functions are now curves, the downward shift of the EoM plane results in a truncation of the accessible speed and displacement range. The extent of this truncation depends on the magnitude of the reduced parasitic force, 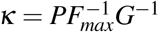, as illustrated by the FL and FV planes, and the EoM plane for *κ* = 0.3. (**C**) The net effect of a constant parasitic force is a decrease in both the maximum relative speed and the effective inertia. (**D**) The magnitude of these changes can be described by proportionality constants, *S*_*i*_, which depend on *κ* and may thus serve as sensitivity indices which quantify the importance of external forces (see eqs. 11a-12): For *κ <* 0.005, *κ <* 0.01 and *κ <* 0.05, all *S*_*i*_ ≥ 0.95, i. e. Γ_*κ*_, Hi_*κ*_ and Bo_*κ*_ are within 5% of Γ, *v*_Hi_, and *v*_Bo_, respectively—the parasitic force may be neglected. The asymptote at 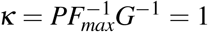 is a force limit to muscle-driven motion. *En route* to this asymptote, all *S*_*i*_ decline steeply. For typical FL and FV relationships, the Hill-number is more sensitive to parasitic forces than the Borelli-number, as the muscle force declines more sharply with strain rate than with muscle strain.

For a Hill-muscle, the influence of parasitic forces is more complex, which I illustrate by returning to the notion of the EoM landscape. For a constant force muscle, the presence of a constant parasitic force manifests itself in a downward shift of the EoM plane (Fig. 3 A). The net force is reduced, but the maximum possible relative speed and displacement remain unchanged, because the muscle force is independent of muscle strain and strain rate; the relative FL and FV functions are equal and constant. For a Hill-muscle, the FL and FV functions are curves. As a consequence of the downward shift of the normalised net force, these curves may now intersect with the zeroplane at a shortening velocity smaller than 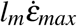, and at a finite displacement smaller than *l*_*m*_*ε*_*max*_; parasitic forces truncate the EoM landscape [Fig. 3 B and 30, 31, 47–51]. The extent of this truncation can, in principle, be quantified through an exercise in equilibrium mechanics [see 30, 47, 50, 51]: the maximum relative displacement over which the muscle can accelerate, and the maximum speed it can contract with, are reached when the net force is zero, i. e. when the muscle force balances the parasitic force. For constant *P*, analytical evaluation of the fraction of the strain rate and strain that is accessible is straightforward: each fraction corresponds to the point on the FL and FV curves at which the muscle force is equal to *P*. Setting *P* equal to the FV and FL functions defined by eqs. 5 and 6 yields:

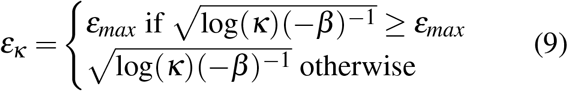

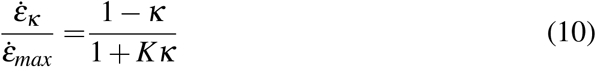

where I introduced the reduced parasitic force, 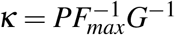. Because the parasitic force truncates both the strain rate and strain range, both the Hill-and Borelli-limit are altered, and follow from combination of eqs. 9 and 10 with eqs.1 and 2, respectively:

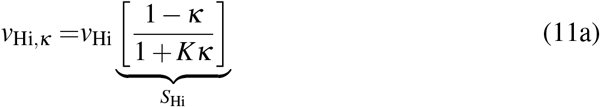

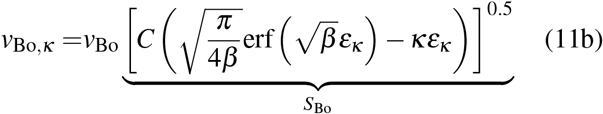

Here, erf is the error function, *C* is a constant that depends on *β* and *ε*_*max*_ (see eq. S25 in the SI), and *S*_*i*_ are sensitivity indices which quantify the importance of the parasitic force (see below and Fig.3 D). Both the Hill- and the Borelli-limit now approach an asymptote of zero speed at a critical reduced parasitic force *κ* = 1; a “force limit” to muscle-driven motion. Close to this limit, both the Hill- and the Borelli-limit drop rapidly (see Figs. 3 C-D). The effective inertia follows immediately as the squared ratio of the Hill- and the Borelli-number:

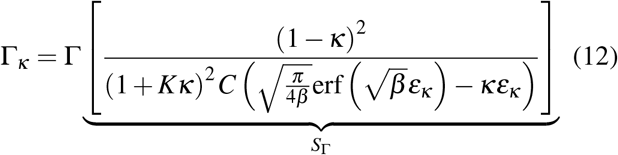

For *G* = 4, and *β* = 6.5, eq. 12 is a decreasing function of *κ* (see Figs. 3 C & D).

The integration of parasitic forces into the definition of the effective inertia, and the visualisation of its effect via the EoM landscape, are conceptually simple. But even for the trivial case of a constant parasitic force, the increase in mathematical complexity is noticeable. Indeed, analytical evaluation of the Hill- and the Borelli-number in the presence of parasitic forces will only rarely be possible, first because of the complex form of the FV and FL relationships, and second because parasitic forces may depend on speed (viscous dissipation), or displacement (structural dissipation), resulting in non-trivial interactions with muscle contraction dynamics. A thorough evaluation of the above results, and their generalisation to characteristic non-constant forces will have to await further work. However, from the cursory analysis presented here emerge two points of note.

First, for a given reduced parasitic force, Γ_*κ*_, *v*_Bo,*κ*_ and *v*_Hi,*κ*_ are constant multiples of Γ, *v*_Hi_ and *v*_Bo_. The proportionality constants, *S*_*i*_, which link both definitions (see bracketed terms in eqs. 11a-12), are thus suitable indices for the sensitivity to parasitic forces: For *κ <* 0.005, *κ <* 0.01 and *κ <* 0.05, all *S*_*i*_ ≥ 0.95 (see Fig 3 D). In other words, Γ_*κ*_, *v*_Hi,*κ*_ and *v*_Bo,*κ*_ are within 5% of Γ, Hi, and Bo, respectively, and the parasitic force may be neglected in leading order analyses. To illustrative the utility of this simple analysis, I address briefly a long-standing observation in comparative biomechanics: small animals are seemingly untroubled by the presence of gravity, and only change locomotor speed slightly, if at all, between running on horizontal or vertical surfaces. In contrast, larger animals generally slow down significantly when moving up inclines [52–55]. The relevant parasitic force is the gravitational force, *F*_*g*_ = *mg*, with an associated reduced parasitic force 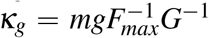, and a ratio between kinetic and parasitic energy — the Froude number of the contraction — 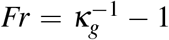. An upper bound *κ*_*g*_ ≤ 0.05*m*^1/3^ kg^−1/3^, where *m* is the body mass in kilogram, was estimated from published data by Alexander [56]. Thus, animals with a body mass *m <* 1 g, *m <* 10 g and *m <* 1 kg may be considered ‘gravitationally indifferent’ in terms of their effective inertia, their Hill- and their Borelli-number, respectively, in robust agreement with the scarce experimental data [52–54]. The work done by the muscle of small animals against the gravitational force is negligible, because the reduced parasitic force is small, and the muscle Froude number is large [see also 18].

Second, the role of musculoskeletal anatomy, represented by the ratio between muscle fibre length and skeletal gear ratio, *l*_*m*_*G*^−1^, needs careful reinterpretation. In the absence of parasitic forces, the Borelli-number is agnostic to musculoskeletal anatomy, because all work flows into kinetic energy, irrespective of whether it is done by displacing a large force over a small distance, or a small force over a long distance (see above and eq. 1). Musculoskeletal anatomy does however control the speed in the Hill-limit, which is maximal for maximum val-ues of *l*_*m*_*G*^−1^ (see eq. 2). In the presence of parasitic forces, both results change fundamentally (see eqs. 11a-11b). In the Borelli-limit, the split of a unit work into force *versus* displacement now matters, because it controls the partitioning into parasitic *versus* kinetic energy; minimising *l*_*m*_*G*^−1^ minimises the re-duced parasitic energy, 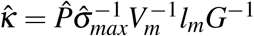. In the Hill-limit, in turn, maximising *l*_*m*_*G*^−1^ will now minimise speed, because it increases the reduced para-sitic force, 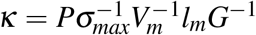, so amplifying the truncation of the accessible contractile speed range [48, 50, the analogous result holds for the truncation of the strain range in the Borelli-limit]. Thus, in both the Hill- and the Borelli-limit, increasing the gear ratio or decreasing the muscle fibre length at constant muscle volume may now increase not only the net force, but also the maximum possible velocity and displacement, in noteworthy contrast to their canonical interpretation as parameters which control putative force-velocity or force-displacement tradeoffs [see above and 57–59, for a recent controversial discussion of this topic]. This force-velocity trade-off is now more complex: if *l*_*m*_*G*^−1^ is too small, the mass can only reach a fraction of its theoretical maximum possible speed, and the muscle only has access to a fraction of its work density; if it is too large, in turn, muscle performance is reduced by parasitic losses instead (see Fig. 3 C–D). The optimum anatomy thus likely corresponds to intermediate val-ues of *l*_*m*_*G*^−1^ which result in effective inertias close to the transition between the Hill- and the Borelli-limit; around this transition, the absolute speed, and relative effective work and effective power density are all close to their theoretical maximum (see Fig. 1 & 2). Realising this optimum will require a different anatomy for a physiologically identical muscle contracting against varying parasitic force (see Fig. 3 C and D). Some reduced parasitic forces, for example those due to gravity, will vary systematically with an-imal size, *κ*_*g*_ ∝ *m*^1/3^ (assuming geometric similarity). It follows at once that there now exists an incentive to depart from geometric similarity via systematic variation of musculoskeletal anatomy, for example due to posture variation [20, 38], in order to remain close to the putative optimum: a systematic increase of gear ratio with size can attenuate the decrease in the Froude number of the muscle contraction predicted from geometric similarity, and keep a larger fraction of the dynamic muscle performance space accessible.

### A theory of physiological similarity in muscle-driven motion

Muscle is a motor with a complexity to baffle any mind; but its operation is eventually restricted to a space bound by the laws of physics. In the above, I have made an attempt to analyse how these laws interact with some of the key physiological and anatomical idiosyncrasies of muscle, in an effort to delineate fundamental limits to its mechanical performance. From this analysis emerged three key dimensionless parameters — the effective inertia Γ, the reduced parasitic energy 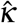, and the reduced parasitic force *κ* — which quantify the relative importance of different physiological and physical constraints on a continuous scale.

The effective inertia, Γ, provides a direct measure of the extent to which the contractile performance of a unit volume of muscle is limited by its strain rate versus strain capacity, and thus of the degree to which it can deliver maximum power and work, respectively. Analogous to how seemingly different problems in fluid dynamics are considered hydrodynamically similar if they occur with equal Reynolds number, problems in muscle-driven motion may be considered *physiologically similar* if they involve equal Γ: the involved muscles will operate with comparable fractions of their maximum strain rate and strain capacity, follow similar contractile trajectories along the EoM landscape, and deliver a similar fraction of their maximum power and work density. A unit volume of muscle has access to its maximum performance space if the musculoskeletal geometry is such that contractions occur with an effective inertia close to unity. For much smaller or much larger Γ, muscle contractions may be considered quasi-instantaneous or quasi-static, and are governed dominantly by the power or work density of muscle, or – equivalently – its FV and FL relationship, respectively. Analytical expressions for the asymptotic maximum speed in these limits are available in form of the Hill-, the Hill-Borelli, and the Borelli-number, which shed light on the distinct relevant “design” features of the musculoskeletal system in each regime.

The reduced parasitic energy and force, 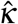 and *κ*, quantify the relative importance of parasitic forces which oppose muscle contraction. Parasitic forces consume muscle work, and truncate the accessible mechanical performance space. As a consequence, the Hill- and the Borelli-number, and thus the effective inertia, gain additional terms which may serve as quantitative indices of sensitivity to parasitic forces. For *κ <* 0.005, parasitic forces may be ignored to first order; for *κ >* 0.005, they alter the dynamics by more than a few percent. Large parasitic forces can decrease both the effective inertia of the contraction, and the maximum speed muscle can impart, resulting in a fundamental change in how musculoskeletal anatomy modulates canonical trade-offs between force and velocity, or force and displacement. It stands to reason that any investigation of muscle performance, and any assessment of musculoskeletal design, ought to explicitly consider Γ, *κ* and 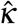of characteristic contractions; ignoring them is to accept considerable risk of erroneous conclusions.

Up to this point, the text may have been heavy in theoretical thought, but arguably light in demonstrated practical consequence. Contractions at different Γ and *κ* may well be governed by rather different physiological and physical constraints, but what range of Γ and *κ* does real muscle find itself in? I note that both Γ and typical *κ* such as *κ*_*g*_ for the gravitational force are size-dependent: Under the parsimonious assumption of geometric similarity, all lengths scale as *L* ∝ *m*^1/3^, all areas as *A* ∝ *m*^2/3^, and all volumes as *V* ∝ *m*. It follows that, for an isogeometric musculoskeletal system moving an isogeometric mass, Γ ∝ *m*^2/3^ and *κ*_*g*_ ∝ *m*^−1/3^. Evolution has thus involuntarily conducted a control experiment in one of the world’s largest laboratories: the animal kingdom. Animals vary by at least 13 or-ders of magnitude in mass, so that, assuming geometric similarity, Γ should vary by a factor about (10^13^)^2/3^ ≈ 10^8^, and 1*/κ*_*g*_ should vary by a factor of (10^13^)^1/3^ ≈ 10^4^. Without allowing for non-trivial departures from geometric similarity, or systematic changes in muscle physiology, it is hard to avoid the conclusion that muscle of animals of different size operates in most substantially different physiological regimes, and with markedly different gravitational sensitivity.

The immediate implications of the influence of Γ and *κ* on muscle performance and of their size-dependence are profound and diverse: characterisation of the maximum intrinsic muscle shortening speed via experiments on isolated muscle fibres is only sound if the contraction occurs in the Hill-limit [see also 35]; extrapolation of such single-fibre experiments to whole muscle or indeed *in vivo* performance can be misleading, as it is typically associated with changes in Γ; musculoskeletal geometry plays a more complex role than its canonical interpretation as a parameter that controls a forcevelocity or forcedisplacement tradeoff would have one believe, so that conclusions from previous comparative analyses of musculoskeletal functional morphology across species may need to be carefully reconsidered [see also 57, 59]; isogeometric animals of different sizes neither have access to the same (maximum) work density, nor to the same (maximum) power density per unit volume of muscle, and are suffering to different extents from parasitic losses to gravitational potential energy — three assertions which challenge the classic scaling theories in animal locomotion [3, 4, 7–9, 17, 21, 22, 33, 60, but see [24]]; and extrapolation of data on extant animals to infer muscle-driven locomotor performance of larger extinct animals may have to explicitly introduce the size-dependence of Γ and *κ*, and assess their influence on the EoM landscape, to avoid physically possible but physiologically prohibited inference. Analyses framed in terms of Γ and *κ* may also provide inspiration for the design of legged robots and in sports biomechanics: electrical motors have characteristic torque-rounds-per-minute relationships, akin to a FV function; many sports may be characterised by specific *κ*, which interacts with the muscle anatomy and physiology of the competing athlete. Some sports may provide dynamic control over *κ*, for example gears in bicycles or oars in rowing, suggesting the possibility of optimal athlete-specific configurations [see 50, for a similar suggestion]. Whether future work verifies or re-jects what at this point are mere hypotheses is immaterial to their potential to advance our understanding of muscle-driven motion, and of the design of musculoskeletal systems.

The insights provided by the theory of physiological similarity should not belie the fact that the list of tasks which need to be completed before it can be considered comprehensive is rather long indeed. It includes the appropriate consideration of activation and deactivation times and the degree of muscle activation [61]; the propagation velocity of muscle excitation [35, 62, 63]; contraction history effects; non-constant parasitic forces and gear ratios [e. g. 23, 64]; muscle pennation [e. g. 65]; series compliance and the associated storage of strain energy in muscle and in tendons [e. g. 16, 64, 66]; multi-segment motion [e. g. 19]; and instances where muscle is lengthened, acts as a break and does negative work [67–69] – to name but a few. I am under no illusion that the simple description of physiological similarity presented here will often be inadequate, and fail to produce a convincing account across the breathtaking diversity of tasks performed by muscle. However, it is my hope that identifying the origin of these shortcomings will push us further in our understanding of the mechanical, physiological, neurological, developmental and phylogenetic constraints that govern the operation of muscle across the animal tree of life.

## Acknowledgments

I am indebted to Ola Birn-Jeffery, Natalie Holt and Angela Kedgley, who all provided most helpful comments on an earlier version of this manuscript. I thank Peter Bishop, Chris Clemente, Taylor Dick, Freddie Püffel, Chris Richards and Jim Usherwood for illuminating exchanges about animal biomechanics throughout the last decade. I am deeply grateful to Natalie Holt who patiently answered a never ending stream of naive questions about all things muscle, and to Chris Clemente for his mentorship. I thank Martin Sprenger for many insightful discussions. This study is part of a project that has received funding from a Human Frontier Science Programme Young Investigator Award (RGY0073/2020) to DL.

## Footnotes

* An exact analysis would need to consider separately the mass of the object and of the muscle. For simplicity, I here lump both into one term. This lumping will introduce an error in calculations where the muscle mass is comparable to the external mass that is moved by the muscle, but this error will only change the result in magnitude, and not in principle, because the distributed mass of the muscle can be taken into account with an “effective mass”-term that is a fixed fraction of the total muscle mass [35].

† For a muscle which generates a constant force throughout the contraction, 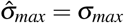. But this is not true for ‘real’ muscle, for which the force varies with both strain and strain rate as discussed in more detail later.

‡ In fact, the power density goes to zero as the strain rate approaches 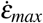, because the Hill-relation is asymptotic, see SI

