## Supplementary Information for "A theory of physiological similarity in muscle-driven motion"

David Labonte

Department of Bioengineering, Imperial College London

### Estimation of the effective inertia of real muscles and musculoskeletal systems

To facilitate estimates of the effective inertia of real muscles and musculoskeletal systems, I re-write eq. 3 in the main manuscript as:

$$\Gamma = \frac{1}{2}\left[\frac{l_m^2}{G^2}\right]\left[\frac{\dot{\varepsilon}_{max}^2}{W_{\rho,max}}\right]\left[\frac{m}{m_m}\right] \qquad \text{(S 1)}$$

Here, $m$ is the mass moved, $m_m$ is the muscle mass, $W_{\rho,max}$ is the maximum work density of muscle, $l_m$ is the muscle fibre length, $G$ is the gear ratio, and $\dot{\varepsilon}_{max}$ is the maximum muscle strain rate. I underline that what follows is an approximate calculation. A (more) exact calculation would account for the movement of the muscle mass separately via continuum mechanics modelling, and distinguish between the muscle mass and the mass that is geared (see the related footnote in the main manuscript).

The advantage of this writing is that $W_{\rho,max}$ is thought to be conserved across vertebrates, with a reasonable value of $W_{\rho,max} = 75$ J kg$^{-1}$ (Alexander, 2003, and eq. S 16 below for a simple model calculation). $\dot{\varepsilon}_{max}$ falls between 1-20 muscle fibre lengths per second for most muscle (Josephson, 1993; Medler, 2002), and a reasonable representative value is $\dot{\varepsilon}_{max} = 5$ muscle fibre lengths per second.[‡] Thus:

$$\Gamma \approx \frac{1}{6}\left[\frac{l_m^2}{G^2}\right]\left[\frac{m}{m_m}\right] \text{meter}^{-2} \qquad \text{(S 2)}$$

For a muscle contracting against itself, $m = m_m$ and $G = 1$, so that $\Gamma \approx 1/6\, l_m^2$ meter$^{-2}$. For a more complex musculoskeletal system, estimates for (i) the ratio between driven mass and muscle mass; (ii) a representative muscle fibre length; and (iii) a representative gear ratio are required. Two of the best studied systems are the hind- and forelimbs of tetrapods, for which large comparative datasets exist. I here focus on the hindlimb, but the consulted references also provide forelimb data; the difference is small. I use a intermediate value of $G \approx 0.5$ (Biewener, 1989), $m/m_m^{-1} \approx 20$, and $l_m \approx 0.027\, m^{0.3}$ meters kg$^{-0.3}$,[‡] obtained from log10-regression on data of hindlimb muscles of 31 mammals and reptile species covering four orders of magnitude in body mass (Bishop et al., 2021). The effective inertia of a generalised tetrapod hindlimb contracting to move the body mass then follows as:

$$\Gamma \approx \frac{1}{100} m^{0.6}\ \text{kg}^{-0.6} \qquad \text{(S 3)}$$

Both results retain significant uncertainty, in particular because $\dot{\varepsilon}_{max}$ and $G$ enter the calculation as a square, and in reality vary to some extent across animals. Thus, these estimates are likely no better than an order of magnitude for any specific muscle or any specific tetrapod. However, due to the strong dependence on fibre length and body mass, comparisons across muscles or tetrapods of disparate size are probably as good as the implicit assumption of geometric similarity. In the absence of robust measurements of all relevant parameters for specific systems, the effective inertia is thus mostly useful in scaling analyses, in keeping with the general idea of similarity arguments.

### The Hill-Borelli-number

The starting point of the derivation is the equation of motion for a general Hill-muscle, which has a characteristic normalised force-velocity (FV, $f(\dot{\varepsilon}_{rel})$) and force-length relationship (FL, $f(\varepsilon)$); $\dot{\varepsilon}_{rel} = \dot{\varepsilon}\dot{\varepsilon}_{max}^{-1}$ is the strain rate normalised with its maximum, and

[‡] The literature is likely biased towards an interest in fast muscle, and maximum strain rates above 10 muscle lengths per second are probably exceptionally large.

[‡] The characteristic muscle fibre length used here is the fibre length of a hypothetical muscle which has a volume and physiological cross-sectional area equal to the sum of all relevant limb muscles. See the original reference for details.

$\varepsilon = \Delta l_m l_m^{-1}$, is the muscle strain; as in the main manuscript, I assume that the optimal length at which the muscle produces maximum force is equal to the fibre resting length, $l_m = l_{opt}$, for simplicity. Both functions are independent, and modulate the muscle force, $F_m$ to a fraction of its maximum, $F_{max}$, so that one may write:

$$a = \frac{F}{m} = \frac{F_m G}{m} = \frac{F_{max} G}{m} f(\dot{\varepsilon}_{rel}) f(\varepsilon) \qquad \text{(S 4)}$$

Here, $G$ is the gear ratio, and $m$ is the mass the muscle is contracting against. The main problem in evaluating this equation is that the muscle strain, $\varepsilon$, and strain rate, $\dot{\varepsilon}$, are coupled to the displacement, $\delta$, and the speed, $v$, of the mass via $\delta = l_m \varepsilon G^{-1}$ and $v = l_m \dot{\varepsilon} G^{-1}$.[‡] Equation S 4 is thus a non-linear differential equation for all but trivial FL and FV relationships. A closed-form solution can however be obtained in some cases, by re-writing eq. S 4 as a path integral, which permits separation of variables:

$$\begin{aligned} \frac{dv}{dt} d\delta &= \frac{F_{max} G}{m} f(\dot{\varepsilon}_{rel}) f(\varepsilon) d\delta \\ \int v \frac{1}{f(\dot{\varepsilon}_{rel})} dv &= \int \frac{F_{max} G}{m} f(\varepsilon) d\delta \end{aligned} \qquad \text{(S 5)}$$

where I multiplied both sides of the equation with $d\delta$, and used $a = dv/dt$, and $d\delta/dt = v$. For trivial FL and FV relationships, i. e. $f(\varepsilon) = f(\dot{\varepsilon}_{rel}) = 1$, eq. S 5 is the familiar Work-Energy-Theorem for a constant net force. The Hill-limit, $v_{\mathrm{Hi}}$, the Borelli-limit, $v_{\mathrm{Bo}}$, and the effective inertia, $\Gamma$, then follow from evaluation of each integral for the maximum possible upper integration boundary:

$$\begin{aligned} \int_0^{v_{max}} m v dv &= \int_0^{\delta_{max}} F_{max} G d\delta \\ \frac{1}{2} m \left( \frac{l_m \dot{\varepsilon}_{max}}{G} \right)^2 &= \hat{F}_{max} l_m \varepsilon_{max} \\ \Gamma = \frac{1}{2} \left[ \frac{l_m^2}{G^2} \right] \left[ \frac{\dot{\varepsilon}_{max}^2}{\hat{\sigma}_{max} \varepsilon_{max}} \right] \left[ \frac{m}{V_m} \right] &= \frac{E_{max}}{W_{max}} \end{aligned} \qquad \text{(S 6)}$$

where $\hat{F}_{max}$ is strictly the average force exerted as the mass is displaced by $\delta_{max}$, and care is required to correctly account for gearing. $\Gamma$ may thus also be interpreted as the ratio between the kinetic energy associated with a contraction at maximum strain rate, and the work done in a contraction to maximum strain. This writing as the ratio of two integrals may also be used to illuminate a more mathematical interpretation of $\Gamma$ for a constant force muscle: its magnitude determines whether the velocity or the path integral have a fixed vs variable upper integration limit, i. e. whether the strain or the strain rate are the state variable vs. fixed parameter, respectively.

For non-trivial FL and FV relationships, life is less kind. For the particular choices introduced in the main manuscript (eqs. 5 & 6), one obtains:

$$\int v \left[ \frac{1 + vKv_{\mathrm{Hi}}^{-1}}{1 - vv_{\mathrm{Hi}}^{-1}} \right] dv = \int \frac{F_{max} G}{m} \exp\left[ -\beta \left( \delta \frac{G}{l_m} \right)^2 \right] d\delta \quad \text{(S 7)}$$

where I introduced the Hill-limit, $v_{\mathrm{Hi}} = l_m \dot{\varepsilon}_{max} G^{-1}$ (see eq. 2 in the main manuscript). This writing can be used to illustrate the difference between the Hill-, Borelli- and Hill-Borelli-limit, with due reference to the earlier comments on the result for a constant force muscle: the muscle is Hill-limited when the upper integration limit for the speed integral is fixed, and the strain is to be determined; it is Hill-Borelli-limited when the upper integration limit for the path integral is fixed, and the strain rate is to be determined; and it is Borelli-limited only as $\Gamma \rightarrow \infty$ so that $\dot{\varepsilon}_{rel} \rightarrow 0$, and $f(\dot{\varepsilon}_{rel}) \rightarrow 1$ throughout the contraction (see main manuscript).

For a constant force muscle, the relevant limit and thus the fixed integration boundary are determined by the magnitude of the effective inertia. For a Hill-muscle, there is a mathematical subtlety: the Hill-relation, which defines the FV relationship, is asymptotic; the maximum strain rate is only reached in the limit of an infinitely long contraction. Notationally, a Hill-muscle thus always reaches $\varepsilon_{max}$ before it reaches $\dot{\varepsilon}_{max}$, and the integration limit of the path integral is thus always fixed. The advantage of this peculiar feature of the Hill-relation is that evaluation of eq. S 7 yields a speed which is the correct asymptotic limit for any value of $\Gamma$. However, this is clearly a mathematical and not a physiological or physical result—no muscle contracts for an infinite time, $\dot{\varepsilon}_{max}$ as defined by the Hill-relation is a purely mathematical construct, and the Hill-limit is thus exact only in the unphysical limit of an infinitely long contraction. In practise, this problem can be resolved by defining a cut-off close to $\dot{\varepsilon}_{max}$ as a realistic maximum strain rate; I consider a muscle to be Hill-limited when the strain rate reaches $\dot{\varepsilon}_c = 0.99\dot{\varepsilon}_{max}$, but any other choice

[‡]Provided that in-series elasticity, for example due to tendon-stretch, is absent.

is possible, and does not influence the essence of the results presented in this work.

Solution of the path integral on the right hand-side of eq. S 7 yields the mass-specific work done in a quasi-static contraction, as before, and is thus related to the Borelli-limit via (see eq. 1 in the main manuscript):

$$\int_0^{\delta_{max}} \frac{F_{max} G}{m} \exp\left[-\beta\left(\delta\frac{G}{l_m}\right)^2\right] d\delta = \frac{1}{2} v_{\mathrm{Bo}}^2 \quad \text{(S 8)}$$

A closed-form solution of the integral is provided in eq. S 16 below.

The speed integral also has a closed-form solution for any real value of $K$, but it involves $v$ both in a logarithmic and a square term, so that no explicit writing is possible; it can of course be solved numerically. For the special case $K = 0$, corresponding to a linear FV-relationship, an explicit solution in terms of $v$ exists:

$$\int_0^{v_{\text{Hi-Bo}}} v \left[\frac{1}{1 - v v_{\mathrm{Hi}}^{-1}}\right] dv = \quad \text{(S 9)}$$

$$-\frac{v_{\mathrm{Hi}}^{-1} v_{\text{Hi-Bo}} + \log\left[1 - v_{\mathrm{Hi}}^{-1} v_{\text{Hi-Bo}}\right]}{\left[v_{\mathrm{Hi}}^{-1}\right]^2} \quad \text{(S 10)}$$

As per above, this integral is equal to $1/2 v_{\mathrm{Bo}}^2$, so that the Hill-Borelli-limit follows as:

$$v_{\text{Hi-Bo}} = v_{\mathrm{Hi}}\left[1 + \mathrm{W}\left(-\exp\left(-1 - \frac{1}{2\Gamma}\right)\right)\right] \quad \text{(S 11)}$$

where W is the Lambert W (or ProductLog) function, and I used $\Gamma = \left(v_{\mathrm{Hi}} v_{\mathrm{Bo}}^{-1}\right)^2$. From this result, it is straightforward to determine critical cut-off values for the effective inertia below or above which a Hill-muscle is effectively Hill- vs Borelli-limited, respectively. Demanding that the Hill-Borelli-limit is to be within 1% of the Hill- and the Borelli-limit yields critical values of $\Gamma_{crit,\mathrm{Hi}} = 0.14$ and $\Gamma_{crit,\mathrm{Bo}} = 1100$.

### Work and power density of muscle and its relation to the effective inertia

Consider a muscle which generates a constant maximum average force $\hat{F}_{max}$ as it moves through a maximum displacement $\delta_{max} = l_m \varepsilon_{max}$. The maximum work per unit muscle mass reads:

$$\frac{W_{max}}{m_m} = W_{\rho,max} = \frac{\hat{F}_{max} l_m \varepsilon_{max}}{m_m} = \frac{V_m \hat{\sigma}_{max} \varepsilon_{max}}{m_m} = \frac{\hat{\sigma}_{max} \varepsilon_{max}}{\rho} \quad \text{(S 12)}$$

where $l_m$ is the length of the muscle, $V_m$ its volume, and $\rho$ the density of the muscle tissue.

Consider next the same muscle which contracts to its maximum strain rate, $v_{max} = l_m \dot{\varepsilon}_{max}$. The maximum average power per unit mass reads:

$$\frac{\hat{P}_{max}}{m_m} = \hat{P}_{\rho,max} = \frac{\hat{F}_{max} l_m \dot{\varepsilon}_{max}}{2 m_m} = \frac{V_m \hat{\sigma}_{max} \dot{\varepsilon}_{max}}{2 m_m} = \frac{\hat{\sigma}_{max} \dot{\varepsilon}_{max}}{2\rho} \quad \text{(S 13)}$$

where the hat now indicates a quantity averaged over the time, and the factor 1/2 arises because the average power is equal to the product between the force and the average speed for a constant force muscle, and $v_{avg} = 1/2 l_m \dot{\varepsilon}_{max}$.

Next, I consider the ratio between the kinetic energy muscle can impart to a mass $m$, and the maximum work that it can do. This ratio may be interpreted as the relative or effective work density of the muscle:

$$W_{\rho,\%} = \frac{m v^2}{2 V_m \hat{\sigma}_{max} \varepsilon_{max}} \quad \text{(S 14)}$$

In the Hill-limit, the maximum speed is $v = v_{\mathrm{Hi}} = l_m \dot{\varepsilon}_{max} G^{-1}$, where $\dot{\varepsilon}_{max}$ is the maximum strain rate; in the Borelli-limit it is $v = v_{\mathrm{Bo}} = \sqrt{2 V_m \hat{\sigma}_{max} \varepsilon_{max} m^{-1}}$. It follows at once that $W_{\rho,\%} = \Gamma$ in the Hill-limit, and $W_{\rho,\%} = 1$ in the Borelli-limit.

In direct analogy, the relative or effective power density is defined as the ratio between the average power the muscle delivers as it accelerates a mass $m$ to a speed $v$, and the maximum average power it could deliver:

$$P_{\rho,\%} = \frac{\hat{F}_{max} G v}{\hat{F}_{max} l_m \dot{\varepsilon}_{max}} = \frac{G v}{l_m \dot{\varepsilon}_{max}} \quad \text{(S 15)}$$

Replacing $v$ with the maximum speed in the Hill- and the Borelli-limit, respectively, immediately yields $P_{\rho,\%} = 1$ in the Hill-limit, and $P_{\rho,\%} = \sqrt{\Gamma^{-1}}$ in the Borelli-limit.

From the above, it follows that muscle operates with maximum power density, but only a fraction $W_{\rho,\%} = \Gamma$ of its work density in the Hill-limit; and with maximum work density, but only a fraction $P_{\rho,\%} = \sqrt{\Gamma^{-1}}$ of its power density in the Borelli-limit. Because $P_{\rho,\%}$ is unity in the Hill-limit, and $W_{\rho,\%}$ is unity in the Borelli-limit, $\Gamma$ and $\sqrt{\Gamma^{-1}}$ also represent the product $W_{\rho,\%}P_{\rho,\%}$ in each limit.

For a Hill-muscle, the force varies with both strain and strain rate, so that the above calculations are more complex. For the FV and FL relationships used in the main manuscript (eqs. 5 & 6), an analytical solution can however be found, by use of the observation that, to first order, contractions that involve maximum power density are governed by the FV function and occur in the Hill-limit, whereas contractions which involve maximum work density occur in the Borelli-limit and are governed by the FL function.

Consider first the work done as the Hill-muscle contracts to a strain $\varepsilon_{max}$ in the Borelli-limit, so that $f(\dot{\varepsilon}_{rel}) \approx 1$. The work done follows as:

$$W_{max} = F_{max} l_m \int_0^{\varepsilon_{max}} \exp\left[-\beta\varepsilon^2\right] d\varepsilon$$
$$= V_m \sigma_{max} \sqrt{\frac{\pi}{4\beta}} \operatorname{erf}\left[\sqrt{\beta}\varepsilon_{max}\right] \qquad \text{(S 16)}$$

where erf is the Gauss error function, and I note that $\hat{\sigma}_{max} = W_{max}(V_m\varepsilon_{max})^{-1}$, and $1/2mv_{\text{Bo}}^2 = W_{max}$ by definition. The maximum work done by Hill-muscle is thus smaller than the maximum work done by a constant force muscle by a factor $\varepsilon_{max}^{-1}\left(\sqrt{\frac{\pi}{4\beta}}\operatorname{erf}\left[\sqrt{\beta}\varepsilon_{max}\right]\right)$. For typical values of $\rho = 1040\,\text{kg}\,\text{m}^{-3}$ (water), $\beta = 6.5$ (Püffel et al., 2023), $\sigma_{max} = 300\,\text{kPa}$ and $\varepsilon_{max} = 0.3$ (Alexander, 2003; Biewener and Patek, 2018), the maximum work density is $W_{\rho,max} \approx 72\,\text{J}\,\text{kg}^{-1}$, comparable to estimates available in the literature (Alexander, 2003).

Consider next the power delivered as a Hill-muscle contracts in the Hill-limit, so that $f(\varepsilon) \approx 1$. The equation of motion reads:

$$\ddot{x} = \frac{F_{max}G}{m}\left[\frac{1 - \frac{\dot{x}G}{v_{max}}}{1 + K\frac{\dot{x}G}{v_{max}}}\right] \qquad \text{(S 17)}$$

Eq. S 17 is a first-order nonlinear ordinary differential equation. For $y(0) = 0$, an analytical solution for the first time integral may be found via the Lambert $W$-function. I will not reproduce this solution here, because it is as lengthy as it is ugly, but instead provide the final result for the time it takes to reach a relative velocity $v_{rel} = \dot{x}Gv_{max}^{-1}$:

$$t_{max} = \frac{m\dot{\varepsilon}_{max}l_m}{F_{max}}\left[(K+1)\log(1 - v_{rel}) + Kv_{rel}\right] \qquad \text{(S 18)}$$

Note that this time diverges as $v_{rel} \to 1$, reflecting the fact that the Hill-equation is asymptotic, and achieves maximum speed only for infinitely long contractions. The maximum average power delivered may be evaluated via $P_{max} = W_{max}t_{max}^{-1}$, where the maximum work is $W_{max} = 1/2(v_{rel}\dot{\varepsilon}_{max}l_m)^2 m$, so that:

$$P_{max} = \frac{v_{rel}^2\dot{\varepsilon}_{max}l_m\hat{F}_{max}}{2\left[(K+1)\log(1 - v_{rel}) + Kv_{rel}\right]} \qquad \text{(S 19)}$$

The maximum power density follows by substituting $F_{max} = \sigma_{max}A_m$, $l_mA_m = V_m$, and dividing by $m_m = \rho V_m$:

$$P_\rho = \frac{\dot{\varepsilon}_{max}\hat{\sigma}_{max}}{2\rho}\left[\frac{v_{rel}^2}{\left[(K+1)\log(1 - v_{rel}) + Kv_{rel}\right]}\right] \qquad \text{(S 20)}$$

The maximum average power density of Hill-muscle is thus also smaller than the maximum average power density of a constant force muscle, by a factor equal to the right-hand term in parentheses. The variation of this term for $0 < v_{rel} < 1$ is not trivial to assess analytically, and, to the best of my judgement, finding the local maxima requires numerical solution. For $\rho = 1040\,\text{kg}\,\text{m}^{-3}$, and typical values of $\dot{\varepsilon}_{max} = 10\,\text{L}\,\text{s}^{-1}$, $\sigma_{max} = 300\,\text{kPa}$ and $K = 4$ (Alexander, 2003), we find a maximum average power density of $P_{\rho,max} = 250\,\text{W}\,\text{kg}^{-1}$ for a contraction to $v_{rel} \approx 0.48 \approx 0.5$, well within the range reported in the literature (Alexander, 2003; Biewener and Patek, 2018). For a contraction to $v_{rel} = 0.99$, the power density drops to $P_\rho = 75\,\text{W}\,\text{kg}^{-1}$.

### The Hill- and Borelli-number for a Hill-muscle in the presence of parasitic forces

Parasitic forces change the Hill- and the Borelli-number, because they truncate the EoM landscape. The Hill-number in the presence of a constant parasitic force $P$ is straightforward to derive, as demonstrated in the main manuscript (eq. 11a). Evaluation of the Borelli-number is more challenging, as it involves both the reduced parasitic energy, and the reduced parasitic force. For a Hill-muscle contracting in the Borelli-limit, the reduced parasitic energy can be evaluated exactly, though the solution is not exactly pretty. The average muscle force, $\hat{F}_m$, may be written as the work divided by the displacement:

$$\hat{F}_m = \frac{W}{l_m \varepsilon_\kappa} = \frac{F_{max}\sqrt{\frac{\pi}{4\beta}}\text{erf}\left[\sqrt{\beta}\varepsilon_\kappa\right]}{\varepsilon_\kappa} \tag{S 21}$$

where $\varepsilon_\kappa$ is the truncated maximum strain, evaluated in the main manuscript (eq. 10). The reduced parasitic energy, $\hat{\kappa}$, follows at once as:

$$\hat{\kappa} = \frac{P}{\hat{F}_m G} = \frac{P\varepsilon_\kappa}{F_{max}G\sqrt{\frac{\pi}{4\beta}}\text{erf}\left[\sqrt{\beta}\varepsilon_\kappa\right]} \tag{S 22}$$

The work that flows into kinetic energy then reads:

$$W_{kin,\kappa} = \left[F_{max} l_m \sqrt{\frac{\pi}{4\beta}}\text{erf}\left[\sqrt{\beta}\varepsilon_\kappa\right]\right][1-\hat{\kappa}] \tag{S 23}$$

$$= F_{max} l_m \left[\sqrt{\frac{\pi}{4\beta}}\text{erf}\left[\sqrt{\beta}\varepsilon_\kappa\right] - \kappa\varepsilon_\kappa\right] \tag{S 24}$$

where I used the result from eq. S 22 in the last step. Division by the maximum work that the muscle can do in the absence of parasitic forces yields:

$$\frac{W_{kin,\kappa}}{W_{kin}} = \frac{\sqrt{\frac{\pi}{4\beta}}\text{erf}\left[\sqrt{\beta}\varepsilon_\kappa\right] - \kappa\varepsilon_\kappa}{\sqrt{\frac{\pi}{4\beta}}\text{erf}\left[\sqrt{\beta}\varepsilon_{max}\right]} \tag{S 25}$$

I define $C = \left[\sqrt{\frac{\pi}{4\beta}}\text{erf}\left[\sqrt{\beta}\varepsilon_{max}\right]\right]^{-1}$, which then yields eq. 11b in the main manuscript.
